# DNA methylation mediated downregulation of histone H3 variant H3.3 affects cell proliferation contributing to the development of HCC

**DOI:** 10.1101/2020.05.23.112516

**Authors:** Divya Reddy, Saikat Bhattacharya, Sanket Shah, Mudasir Rashid, Sanjay Gupta

## Abstract

H3.3 variant is a versatile histone important for development and disease. We report a DNA methylation dependent decrease of histone H3 variant H3.3 in hepatocellular carcinoma (HCC) development and an increase in the level of the H3.2 variant. The loss of H3.3 correlates with a decrease in the histone PTMs associated with active transcription. The overexpression of H3.3 and H3.2 did not affect global PTMs and cell physiology, probably owing to the deregulation of specific histone chaperones CAF-1 (for H3.2) and HIRA (for H3.3) that we observed in HCC. Notably, upon P150 (CAF-1 subunit) knockdown in HCC cell lines, a cell cycle arrest in S-phase was observed, possibly due to the decrease in the histone levels necessary for DNA packaging. Furthermore, H3.3 knockdown in a preneoplastic liver cell line led to an increase in cell proliferation and a decreased transcription of tumor suppressor genes, recapitulating the tumor cell phenotype. Importantly, our data suggest that the use of DNA Methyl Transferase (DNMT) and Histone Deacetylase (HDAC) inhibitors to restore the expression of H3.3 and the altered chromatin state for the better clinical management of the disease.

## 1. Introduction

Chromatin assumes distinct conformations that regulate the activity or repression of given genes. The cellular states and responses can be distorted by various genetic, metabolic, and environmental signals that disrupt the chromatin, thereby predisposing individuals to a variety of common diseases. Although cancer is usually considered a genetic disease, chromatin and epigenetic irregularities play crucial roles in tumor initiation as well as its progression [1].

Changes in the conformation of the chromatin states are either achieved by DNA methylation, post translational modification (PTM) of histones, or by changing the biochemical composition of the nucleosomes by the replacement of major histone types with specific histone variants via definite histone chaperones [2]. Expression of certain histone variants have been correlated with tumor malignancies. macroH2A, an H2A histone variant, suppresses cancer progression and acts as a direct transcriptional repressor of CDK8, a cancer driver gene [3]. H2A.Z variant promotes cancer progression via transcriptional regulation in prostate and breast cancers [4,5].

Histone H3.3 is one of the very well-studied variants in the context of development, differentiation and disease [6]. H3.3 deficient mice embryos display reduced levels of open chromatin marks like H4K16Ac, leading to the compaction of the chromatin [7]. Further, H3.3 knockout mice have reduced viability, and the surviving adults are infertile [8]. The deposition of H3.3 by its chaperone HIRA has been shown to affect the nucleosome dynamics and thus gene expression, to govern the cell fate transition process [9]. Together, these findings indicate that H3.3 is involved in establishing a finely balanced equilibrium between the open and condensed chromatin states during various cellular processes.

One-way by which the histone variants and their various specific PTMs can influence the various pathways is by affecting the nucleosome dynamics and thus the chromatin organization. Unlike its canonical counterparts H3.1/H3.2, H3.3 incorporation into the chromatin is cell-cycle independent or replication-independent, and it can be deposited at replication sites when the canonical H3.1/H3.2 deposition is impaired [10]. H3.3 has also been consistently associated with an active state of chromatin. Additionally, cancer specific mutations in H3 variants or broadly on the histones itself, which mainly results in changing the PTM profile, thus influencing the underlying gene expression [6].

Given their fundamental role in shaping the chromatin structure, the histone variants (and their chaperones) likely constitute important biomarkers for diagnosis or prognosis and may even represent therapeutic targets in cancer patients. To date, studies have demonstrated the potential prognostic utility of the histone variants in multiple cancers including those of the lung and the breast [13,14]. Furthermore, the cellular levels of histone variants and also PTMs can also foresee responses to certain chemotherapeutic agents, serving as predictive biomarkers that could impact decisions concerning courses of therapy [15]. However, the main challenge currently is to mechanistically understand how histone variant deregulation can contribute to cancer development and progression.

To understand the role of histone variants in cancer, we have used an NDEA-induced hepatocellular carcinoma (HCC) model system in the Sprague-Dawley rats. On screening for histone pattern changes between the control and tumor tissues, we identified that there is DNA methylation mediated downregulation of the transcription of histone H3 variant H3.3 in HCC and a concomitant increase of H3.2 expression. Impairing the deposition of H3.2 by P150 knockdown not only leads to the arrest of the cells in S-phase of the cell cycle but also increases H3.3 protein levels. Further, we also show that H3.3 occupies promoters of tumor suppressor genes and influences their expression. Our report suggests that there is a co-operative interplay between histone variants, histone chaperones, and their transcriptional regulatory machinery resulting in the stable maintenance of the highly dynamic histone marks required in the deregulated epigenetic landscape found in cancer cells.

## 2. Materials and methods

### 2.1. Animal Handling and Experiments

All the experiments were performed using male Sprague-Dawley rats (spp. *Rattus norvegicus*) after the approval of the Institute Animal Ethics Committee (IAEC# 04/2014), Advanced Centre for Treatment Research and Education in Cancer and the Committee for Control and Supervision on Animals, India standards. The protocol to induce liver carcinogenesis is as described previously [16]. Tissue samples were fixed in formalin and prepared as paraffin-embedded blocks according to standard protocols. The H&E-stained sections were then microscopically evaluated for histopathological alterations to validate normal and HCC samples.

### 2.2. Isolation of histones from liver tissue

Histones were extracted and purified as described earlier [17]. Briefly, liver tissues (1gm) were homogenized in 10 ml lysis buffer (15mM Tris-Cl pH7.5, 60mM KCl, 15mM NaCl, 2mM EDTA, 0.5mM EGTA, 0.34M sucrose, 0.15mM β-mercaptoethanol, 0.15mM spermine and 0.5mM spermidine) with 1X protease inhibitor cocktail and phosphatase inhibitor cocktail. Nuclei were then isolated by sucrose gradient centrifugation. The three volumes of homogenate were layered on top of one volume of 1.8M sucrose. Nuclei pellet obtained by centrifugation at 26,000rpm for 90 min at 4°C was resuspended in 0.2M H2SO4 and incubated for >2 hrs at 4°C with intermittent vortexing. After centrifugation at 16,000rpm for 20 min at 4°C, the histones were precipitated at −20°C for overnight with the addition of four volumes of acetone to the supernatant. Post centrifugations at 16,000rpm for 20 mins at 4°C, pelleted histones were air-dried and suspended in 0.1% β-mercaptoethanol in H2O and stored at −20°C.

### 2.3. Resolution and analysis of histones

The purified histones from serum and tissue were resolved on 18% sodium dodecyl sulfate-polyacrylamide gel electrophoresis (SDS-PAGE) was either stained by silver staining method or transferred to PVDF membrane, probed with site-specific modified histones antibodies, against H4K16Ac (Millipore#07-329), H4K20Me3 (Abcam#9053), γH2AX (Millipore#05-636), H3S10P (Millipore#06-570), H3K27Me3 (Millipore#07-449), H3K9Me3 (Abcam#8898), H3K9Ac (Millipore#07-352), H3K27Ac (Abcam#4729), H3K14Ac (Abcam#52946), H3Ac (Upstate#06-599), H3K4Me3 (Abcam#1012), H3 (Upstate#06-755), H4 (Millipore#07-108) and signals were detected by ECL plus detection kit (Millipore #WBKLS0500). Gel loading equivalence was done by the Silver Staining method.

### 2.4. RP-HPLC

The reversed-phase separation was carried out on a C18 column (1.0×250mm, 5 mm, 300Å; Phenomenex). Mobile phases A and B consisted of water and acetonitrile with 0.05% trifluoroacetic acid, respectively. The flow rate was 0.42 ml/min, and the gradient started at 20% B, increased linearly to 30% B in 2 min, to 35% B in 33 min, 55% B in 120 min and 95% B in 5 min. After washing at 95% B for 10 min, the column was equilibrated at 20% B for 30 min, and a blank was run between each sample injection.

### 2.5. Cell line maintenance and synchronization

The cell lines were maintained in DMEM media (Invitrogen) at 37°C with 5% CO_2_ supplemented with 10% FBS, 100U/ml penicillin, 100mg/ml streptomycin and 2mM L-glutamine (Sigma). Cells were enriched in early G1-phase by serum starvation (0.1% FBS) for 24 hrs. Media supplemented with 10% FBS was used to release the cells from the G1-arrest.

### 2.6. shRNA knockdown and selection

shRNA sequences targeting P150 and H3.3 was cloned into pLKO.puro vector using AgeI and EcoRI enzymes. All the primers and shRNA sequences used in the study are tabulated in Table 1. Post transfection by turbofect (Fermentas), stable clones were selected with 1µg/mL Puromycin (Sigma) for 1 week, the clones obtained were then checked for their expression.

**Table 1:**
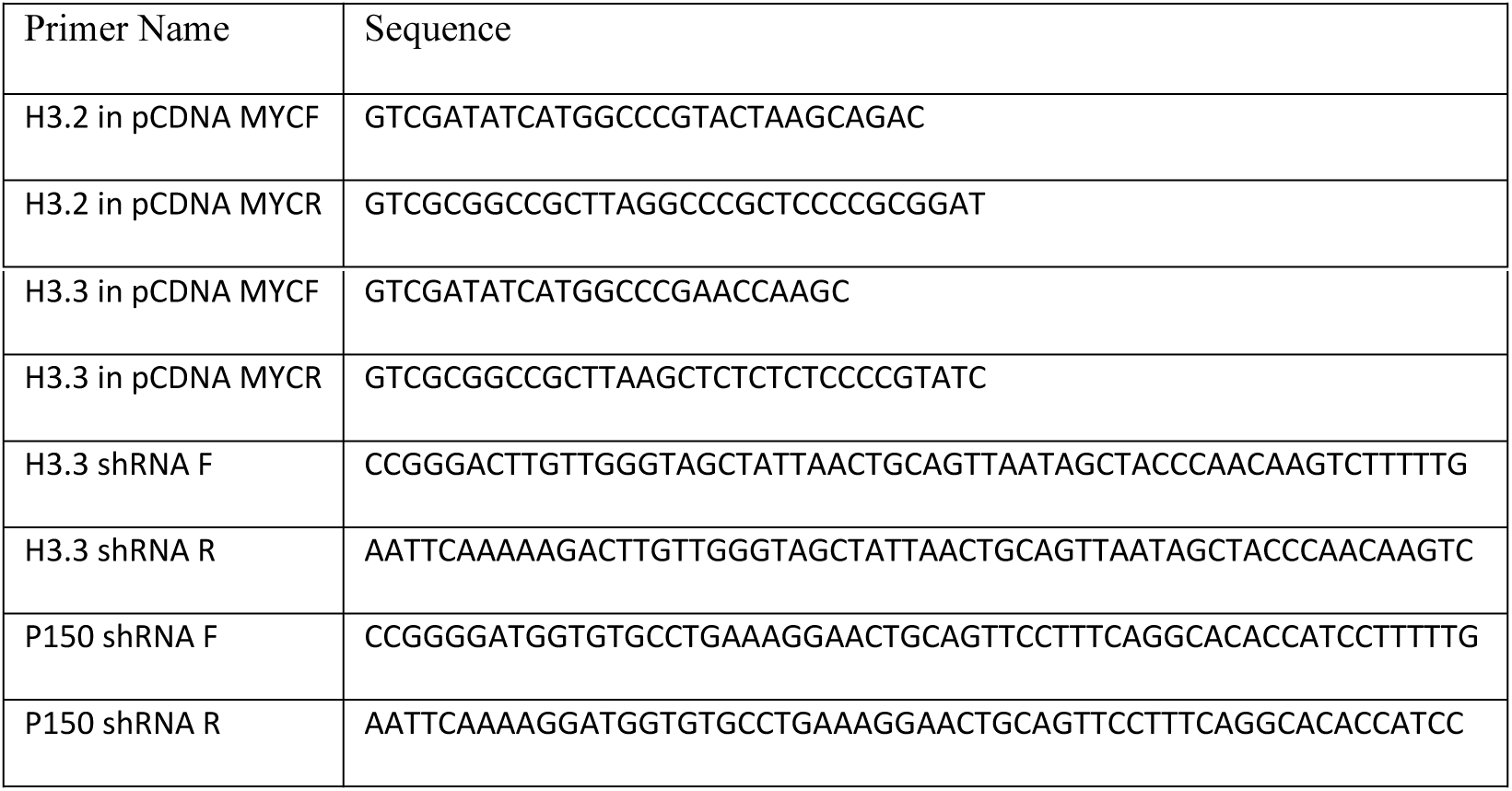
Primers Used for Cloning.

### 2.7. Total RNA isolation and qPCR

Total RNA was extracted from 25 mg of the frozen tumor and control tissues as per manufacturers protocol (Thermo scientific-0731). For cell lines, TRizol method of RNA isolation was employed. Total RNA (1 μg) was used for cDNA synthesis (Fermentas-K1632) using random hexamers. qPCR with SYBR green was done using specific primers tabulated in Table 2. For the histone variants a common primer designed to detect all sequence variations was used.

**Table 2:**
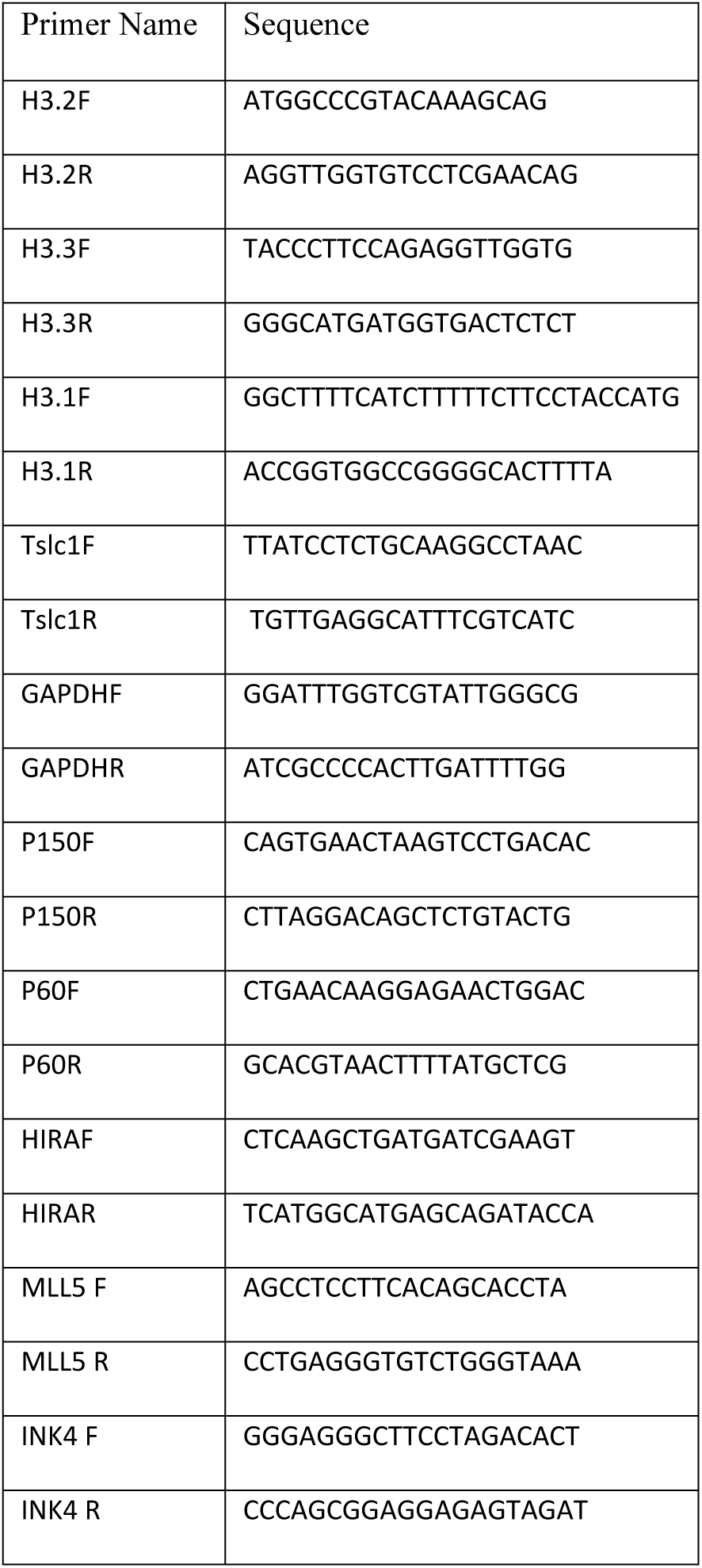
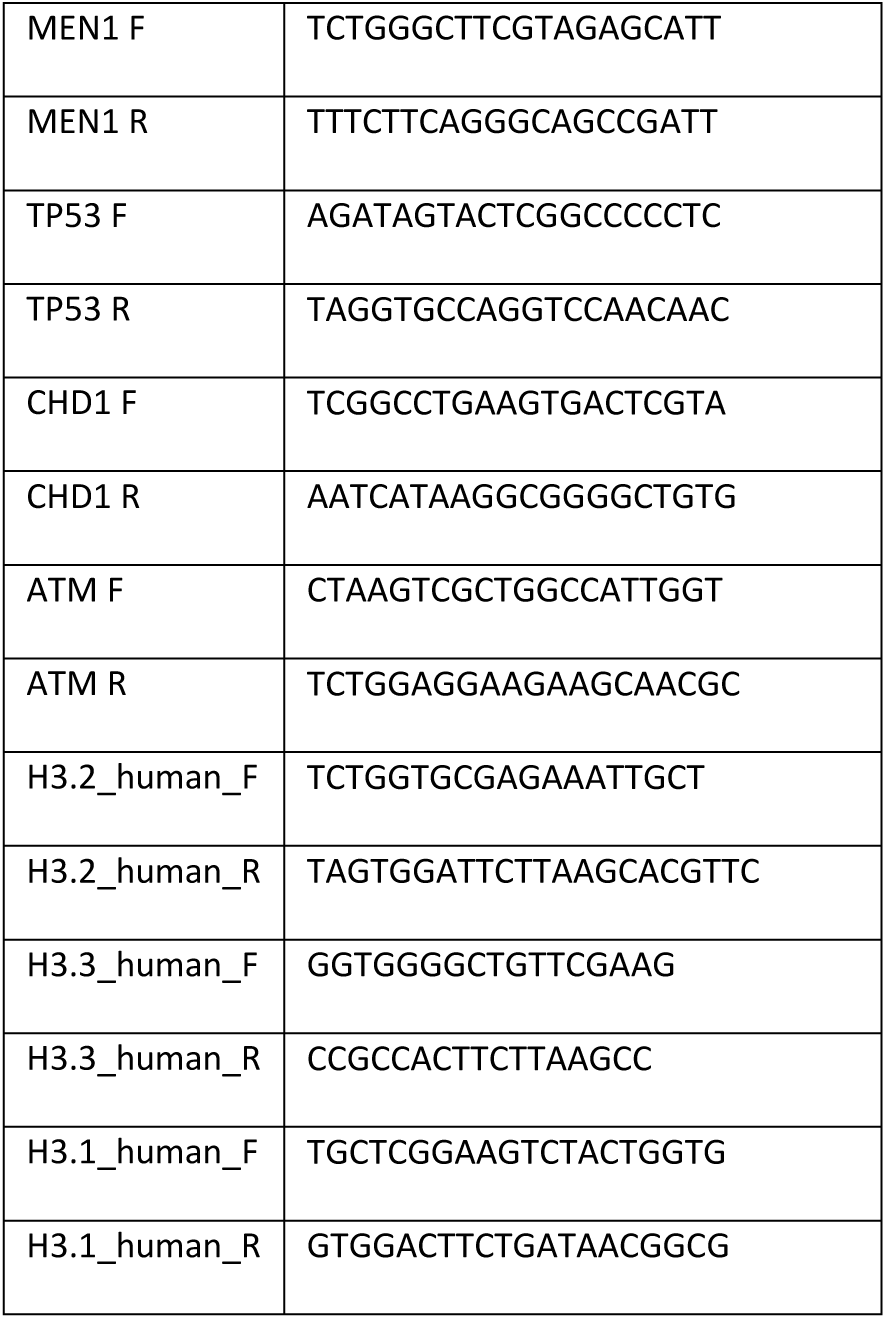
Primers Used for Real time PCR.

### 2.8. Cell cycle analysis

Ethanol fixed cells were washed twice with PBS and suspended in 500µl of PBS with 0.1% Triton X-100 and 100µg/ml of RNaseA followed by incubation at 37°C for 30mins. After incubation, propidium iodide (25µg/ml) was added followed with incubation at 37°C for 30mins. DNA content analysis was carried out in a FACS Calibur flow cytometer (BD Biosciences, USA). Cell cycle analysis was performed using the ModFit software from Verity house.

For Azacytidine and Trichostatin A (TSA) treatment, the cells were cultured in the medium as described above along with 5µM Azacytidine and 10nM TSA for 16hr and 72hr respectively. Post incubation cells were harvested, and RNA was isolated using Trizol reagent. cDNA was synthesized using a random hexamer and gene-specific primers were used for real-time PCR

### 2.9. Methylated DNA immunoprecipitation

Genomic DNA was purified using the sigma gDNA Miniprep kit (G1N350), according to the manufacturer’s instructions. Purified genomic DNA was diluted into a total of 300ml TE buffer and sonicated with a Bioruptor (10 cycles at low power, of 30sec ‘on’ and 30sec ‘off’) to an average size of 300–500bp. An aliquot of sonicated DNA was run on 1% agarose gel to confirm fragment size during each methylated DNA immunoprecipitation (MedIP) procedure. Sonicated DNA (4µg) was denatured by incubation at 95°C for 10min and was then immediately transferred to the ice for 10min. Immunoprecipitation buffer containing 10mM sodium phosphate, 140mM NaCl, and 0.05% Triton X-100 was added to a final volume of 500 µl. For each IP reaction, 2µg of antibody (Methyl cytosine: Diagenode MAb-006-100, Hydroxy methylcytosine: Abcam ab106918) was added and incubated overnight at 4°C with shaking. Five percent of DNA was kept as input. After incubation, 30µl of Dyna Protein G beads (Invitrogen: 10004D) were added and further incubated for 1hr at 4 °C with shaking. Beads were washed thrice with 500 µl of IP buffer. Elution buffer (150µl) containing 50mM Tris-HCl pH 8.0, 10mM EDTA, 1%SDS, 50mM NaHCO_3_ and 20µg proteinase K was added and incubated at 55°C for 3hr. Tubes were applied to a magnetic rack and eluted DNA and input DNA were purified with the Qiaquick PCR purification kit (Qiagen) followed by SYBR Green real-time quantitative PCR to identify methylated regions. PCR measurements were performed in duplicate. The primers designed for H3.2 and H3.3 are part of the CpG sites and CpG island respectively (Table 1). The average cycle thresholds for the technical replicates were calculated to yield one value per primer set for each biological replicate and normalized to input using the formula 2^(**Ct**(input)-{**Ct**(immunoprecipitation))^. Averages and standard deviations of the normalized biological replicate values were plotted in the Figs and used in t-test calculations. The primers used are tabulated in Table 3.

**Table 3:**
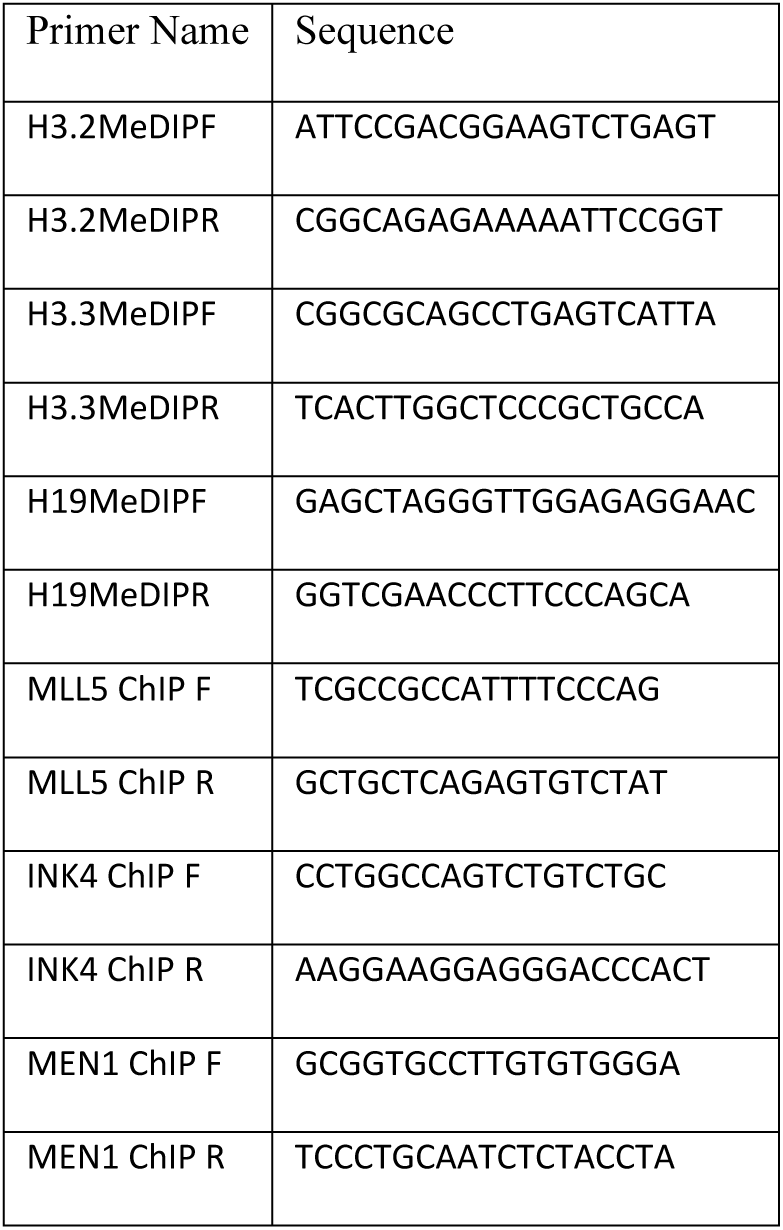
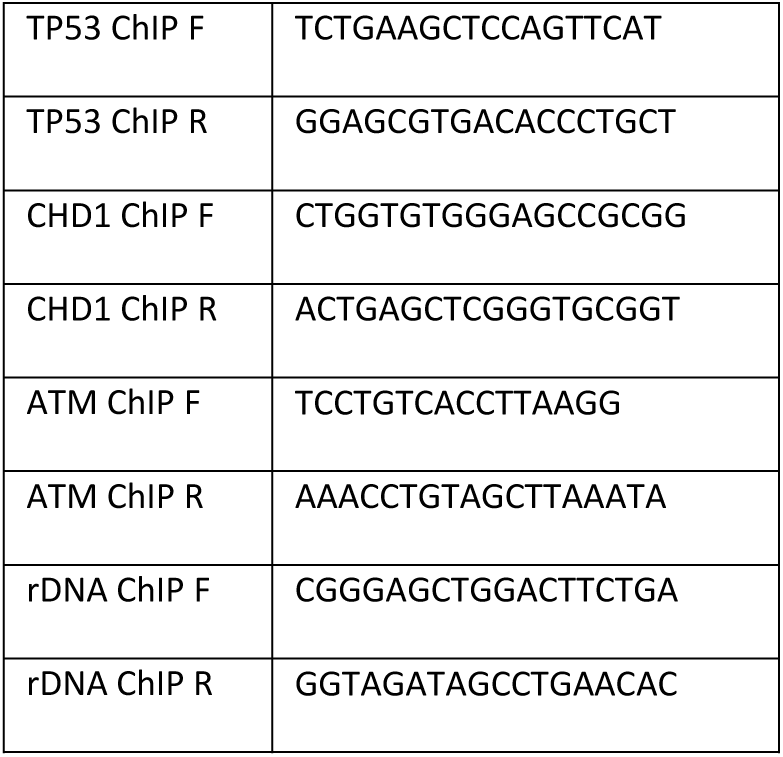
Primers Used for MeDIP and ChIP qPCR.

### 2.10. Chromatin immunoprecipitation

Chromatin immunoprecipitation (ChIP) was carried out as described in the Acetyl-Histone H3 Immunoprecipitation Assay Kit by Millipore. DNA recovered from chromatin immunoprecipitation was analyzed by real-time PCR. The reaction mixture (25 µL) contained 1 µL of the appropriately diluted DNA sample, 0.2 µmol/L primers, and 12.5 µL of IQ SYBR Green Supermix (Bio-Rad). The reaction was subjected to a hot start for 3 min at 95°C and 50 cycles of 95°C, 10 s; 55°C to 65°C, 30 s; and 72°C, 30 s. Melt curve analysis was done to verify a single product species. Percent enrichment in each pulldown was calculated relative to input DNA. The primers used are tabulated in Table 3.

### 2.11. Statistical analysis

All numerical data were expressed as the average of values obtained ± standard deviation (SD). Statistical significance was determined by conducting a unpaired students’ *t*-test or a paired students test where ever applicable.

## 3. Results

### 3.1. Development of HCC is accompanied by a decrease in the level of the histone variant H3.3 and simultaneous increase of H3.2 in chromatin

Sprague-Dawley rats were induced for HCC by administrating the chemical carcinogen NDEA as described earlier [16]. The tissues were collected after sacrificing the rats and the stage of the cancer was confirmed by histopathology. Histones isolated from the control and the tumor liver tissues were resolved using AUT-PAGE which separates proteins based on mass and hydrophobicity [18]. Along with the significant change in the pattern of H2A isoforms H2A.1 and H2A.2, as reported earlier [16], major alterations in the H3 region were also found (Fig 1A). The upper and the lower band were identified as H3.2 and H3.3 respectively by mass spectrometry (Fig 1B). The details of the peptides differentiating the H3 variants have been tabulated in Supplementary Fig 1. <u>R</u>everse-<u>p</u>hase-<u>h</u>igh-<u>p</u>erformance liquid <u>c</u>hromatography (RP-HPLC) done to quantitatively measure the differences in H3 variant profile in control and tumor histones confirmed that H3.2 level was increased and H3.3 was decreased in the tumor tissue (Fig 1C(i)(ii)). Measurement of these transcripts in the tissues revealed a significant upregulation of H3.2 and downregulation of H3.3 in tumor compared to control (Fig 1D(i)). Similar changes were observed in pre-neoplastic CL44 and neoplastic CL38 cells (Fig 1D(ii)). These cell lines were derived from the liver of Sprague Dawley rats post administration of NDEA [19].

**Fig 1:**
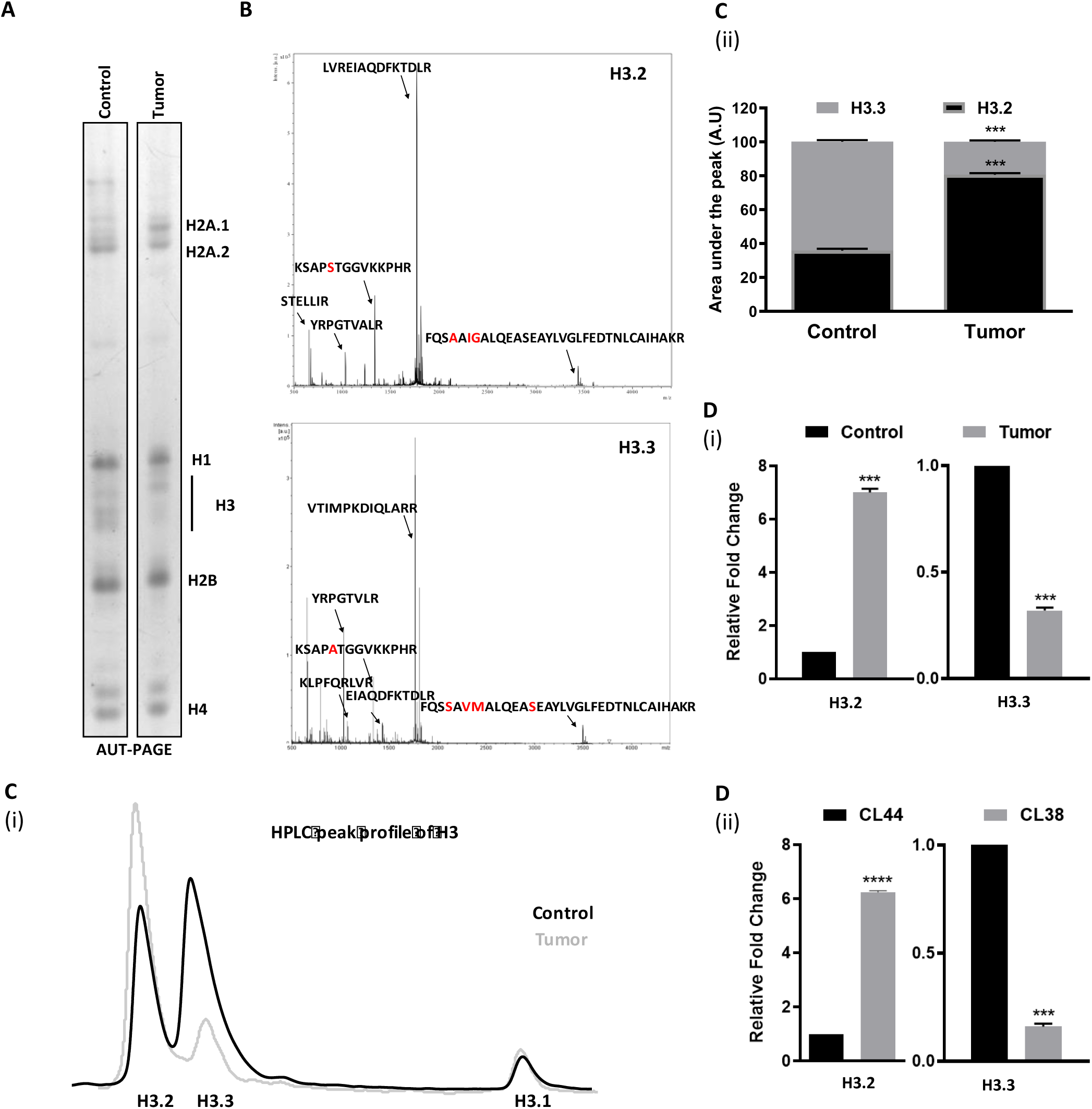
Dysregulation of H3 variants correlates with tumor tissue. (a) Silver stained AUT-PAGE analysis revealed changes in the expression profile of H2A.1, H2A.2 and H3 region. Remaining histones are also marked. (b) Mass spectrometry analysis of the H3 region identified the upper band as H3.2 and lower protein band to be H3.3 respectively. The differential amino acids in the tryptic digested peptides are highlighted in red. (c)(i) Overlay of H3 variant RP-HPLC profile of control and tumor tissues showing upregulation of H3.2 and downregulation of H3.3 in tumor tissue. (ii) The quantitative graph plotted for showing the area under the peak for H3.2 and H3.3. (d) Real-time PCR, done to assess the changes in the transcript levels of H3.2 and H3.3 in (i) tissues and (ii) cell lines.

The transcription of replication-dependent, canonical histones like H3.2 and H3.1 markedly increase upon the entry of cells into S-phase aiding the nucleosome formation on newly synthesized DNA [20]. H3.3 variant, on the other hand, is a replication-independent histone variant and is expressed throughout the cell cycle. As expected, the tumor tissue and the CL38 cells were more in S and G2/M phase compared to normal and CL44 cells respectively (Fig S 2A). We next asked whether only the H3.2 variant is upregulated or the H3.1 variant, which is another replication dependent H3 histone is also upregulated in transformed cells. Strikingly, the expression level of H3.1 was unaltered at the transcript level in both the tissues and cell lines (Fig S 2B). A similar observation was also seen at the protein level (Fig 1C(i)), suggesting that H3.2 is specifically upregulated in HCC.

To understand if these changes in the H3 variant profile is a consequence of increased cell proliferation or is it due to the carcinogenesis phenomenon, a highly proliferating system-regenerating liver post partial hepatectomy (PH) was used. Markers like AFP, Ki67 and Sox 9 were used for validating the regenerating tissue (S3A). Unlike the changes seen in the tumor tissue, PH liver showed elevated levels of all the three H3 variant levels, both at the protein and transcript levels (Fig S3 B, C). This, suggests that the decrease in H3.3 is a cell transformation associated phenomenon and is merely not reflective of the increased cell proliferation observed in cancer. H19 was used as a control, expression of which is known to change in different pathophysiological states of the liver [21]. Intriguingly, even though tumor cells are residing more in S-phase only H3.2 is increased and not the other replication-dependent histone H3.1, indicating that maybe these cells are tuned in such a way to express only H3.2 probably to affect the downstream pathways.

To understand whether these changes in the expression profile of H3 variants, observed in the rat HCC model system is true in human cells also, profiling for H3 variants at the transcript level in 3 different types of cancer cell lines alongside their immortalized or non-transformed counterparts was done. Decreased expression of H3.3 was seen in all the cancer cell lines, HepG2 (liver), MCF7 (breast), and A431 (skin) compared to HHL5 (liver), MCF10A (breast) and HACAT (skin) (Fig 2A (i, ii, iii)). Further, H3.2 was found to be increased in all the tumor cell lines along with H3.1 in skin cancer.

**Fig 2:**
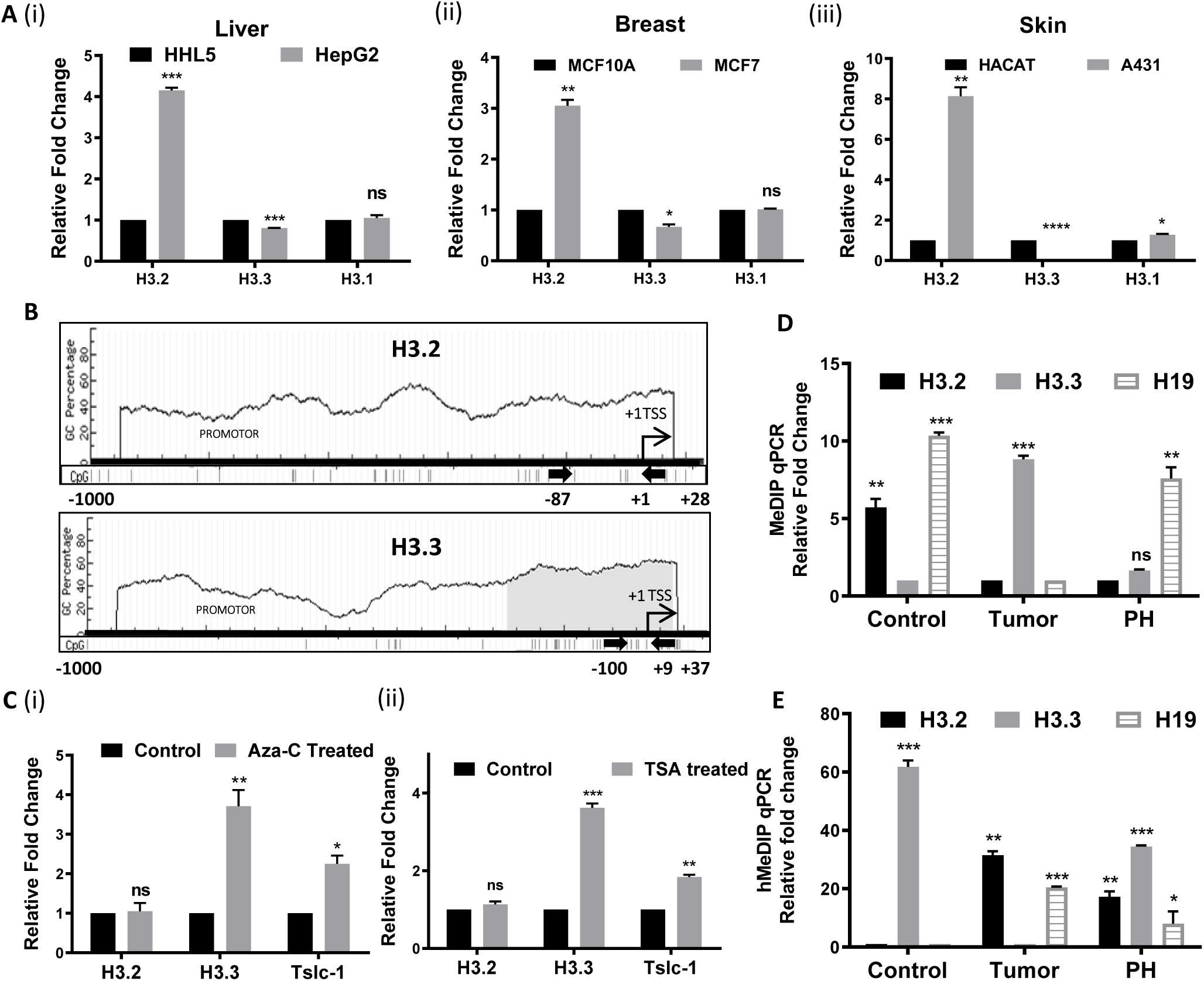
DNA methylation governs expression changes in the H3 variant profile in tumor. (a) Quantitative real-time PCR data showing the relative expression levels of H3.2, H3.3, and H3.1 in Liver (i), Breast (ii), and Skin (iii) cell lines. (b) *In silico* (by Methyl DB) analysis of the promoter elements of H3 variants. +1, Transcriptional Start Site (TSS) has been marked. Arrows indicate the primers used for Methyl Immunoprecipitation analysis. (c) Measurement of relative transcript levels of H3.2 and H3.3 upon treatment with Azacytidine (5 Aza-C) (i) and TSA (ii). Relative enrichment of 5-Methyl Cytosine (e) and 5-hydroxyl-methyl cytosine (f) on promoters of respective histone variants in control tumor and PH tissues.

### 3.2. DNA methylation regulates the expression changes of H3 variants

To understand the reason behind the transcription pattern of H3.2 and H3.3 their promoters were analyzed for any unique sequences. Intriguingly, the H3.3 promoter consisted of a CpG (-CCGG-) island whereas the H3.2 promoter has CpG sites at its predicted transcriptional start site (Fig 2C). DNA methylation and histone modifications are two major epigenetic modulators that are known to regulate gene expression. Increased methylation of CpG islands or sites at the 5’ end of a gene is associated with gene repression. This occurs probably due to inhibition of transcription factor binding directly or by the recruitment of transcription repressors like HDAC’s by methylcytosine binding proteins. We hypothesized that DNA methylation might regulate H3 variant expression changes, to this end, we monitored expression status of H3 variants after treatment of CL38, a neoplastic cell line with small molecule inhibitors of DNMT and HDAC, 5’-Azacytidine (Aza-C) and Trichostatin A (TSA) respectively. Indeed, treatment with both the inhibitors led to the increase in H3.3 with no significant change in H3.2 expression (Fig 2C(i) and (ii)). Tslc-1 was used as a positive control for the inhibitor treatments, as its expression is controlled by DNA methylation [22].

To further understand if indeed DNA methylation directly regulates the H3 variant expression, Methyl DNA immunoprecipitation (MeDIP) following qPCR was performed with control, tumor and PH tissues. This technique employs antibodies against 5-methyl cytosine (5-mC) and hence can be used quantitatively to measure DNA methylations levels on the desired gene. Interestingly, 5-mC, which is a transcriptional inactive mark was found to be higher in tumor tissue in comparison to control on the H3.3 promoter and less on H3.2 promoter. DNA hypomethylation of the H3.2 promoter was also seen in PH tissue (Fig 2D). Methyl cytosine is known to be actively demethylated to produce a series of intermediary products like 5-hydroxymethylcytosine (5-hmC). This is a stable epigenetic mark that has been associated with active gene transcription. Hence, using hydroxyl methyl cytosine antibodies to perform 5-hydroxyl methyl cytosine immunoprecipitation (MeHdIP) analysis is a good estimate for actively expressing genes. MeHdIP analysis of H3 promoters corroborated with the expression status of the variants, high 5-hmC was seen on genes actively expressed - H3.2 in tumor tissue and H3.3 in control tissue (Fig 2E). H19 promoter analysis was used as a positive control for MeDIP, as its gene expression is known to be governed by DNA methylation [23].

Thus, indeed DNA methylation is a dynamic player governing the expression pattern changes of H3.2 and H3.3 in HCC. Interestingly, the changes in the methylation profile on H3 promoters correlates with the hallmark of cancer that is CpG island hypermethylation and global hypomethylation [24]. Further, these changes seem to be a cancer specific phenomenon and not a general aspect associated with cell proliferation, as these are not observed in highly proliferating liver cells after partial hepatectomy.

### 3.3. Specific histone H3 variant levels correlate with the H3 PTM modifications

Histone H3.3 is generally associated with the transcriptionally active chromatin [25]. The downregulation of H3.3 in HCC prompted us to investigate the levels of histone PTMs that are associated with eu- or hetero-chromatin. There was a decrease in the pan H3 acetyl mark in tumor and CL38 cells in comparison to the normal liver tissue and CL44 (Fig 3A). Interestingly, all the histone acetylation marks associated with gene activation like H3K27Ac, H3K14Ac and H3K9Ac were also found to be lower along with H3K4Me3. Further, there was also an enrichment of repressive marks like H3K9Me3 and H3K27Me3 in the tumor cells (Fig 3A). A slight decrease in H4K16Ac, well-established histone PTM hallmark of cancer was also observed.

**Fig 3:**
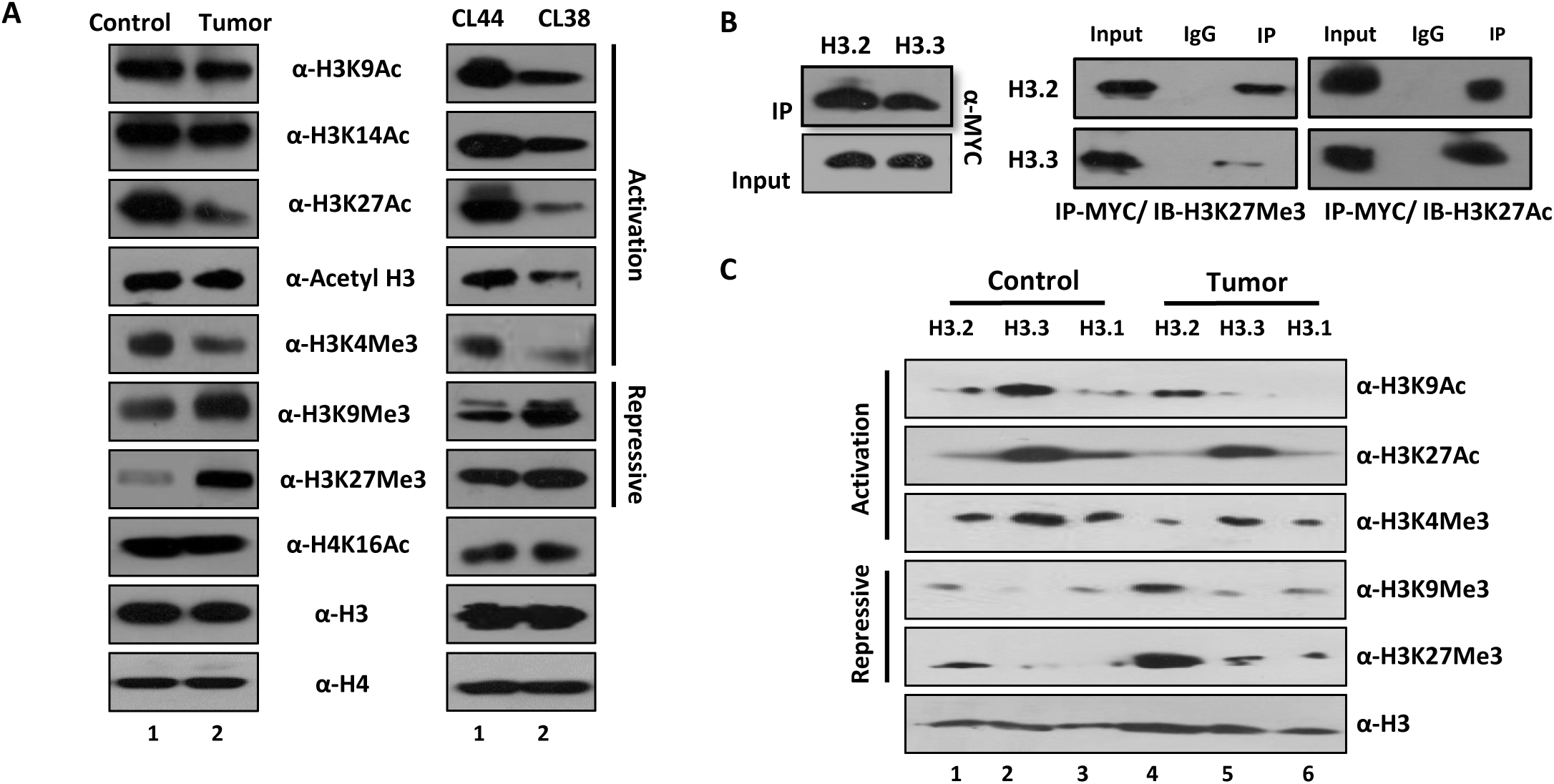
H3.3 is enriched with active histone PTM’s. (a) Western blotting of histones isolated from tumor and control tissues and cell lines with site-specific histone modifications. H3 and H4 probing were used as a loading control. Active and repressive marks are denoted. (b)(i) Mononucleosomal Immunoprecipitation of H3.2 and H3.3 with MYC followed by western blotting with H3K27Me3 (Repressive mark) and H3K27Ac (Active mark) (b)(ii) Monomeric histones of H3 variants separated by HPLC were probed for respective active and repressive histone PTMs for both control and tumor. Western for H3 was used as a loading control.

To know whether the loss of activation marks and gain of repressive marks is due to the changes in H3.3 and H3.2 profile, respectively in tumor tissues and cell lines, two approaches were used. Firstly, ectopically expressed MYC-tagged H3 variants were immunoprecipitated from CL44 cells and probed for H3K27Me3 (inactive mark) and H3K27Ac (active mark). The result suggests that indeed H3.3 is enriched with active mark and H3.2 with inactive mark (Fig 3B). In the second approach, with the help of RP-HPLC, H3.2, H3.3 and H3.1 fraction were separately collected from both control and tumor tissue and probed with active and inactive PTM marks. The loss of the active marks was predominantly seen from H3.3 and the gain of repressive marks on H3.2 (Fig 3C). However, H3.1 also showed a marginal decrease in the active and increase in inactive marks. Nonetheless, the changes in the PTM profile can be majorly attributed to the changes in the variant profile.

### 3.4. Histone H3 chaperone levels are altered in HCC

We showed that DNA methylation mediates a balance of expression of H3 variants, H3.2 and H3.3 in cancer. Next, we postulated that any perturbation in this balance in normal cells may lead to similar changes observed in cancer. To assess this, MYC-tagged H3.2 and H3.3 variants were overexpressed, in CL44 cells. However, this did not lead to any effect on proliferation of cells or any changes in the global histone PTM profile even though these variants are incorporated into chromatin. (Fig S4). This led us to conclude that although the ectopically expressed H3 variants are incorporated into chromatin, it is only exchanged with the respective H3 variant with no overall enrichment.

This is quite possible as H3.2 and H3.3 variants have their dedicated histone chaperones CAF1 and HIRA, respectively [26]. These chaperones are responsible for the recruitment and incorporation of these two variants into their specific genomic loci. The expression levels of the H3 chaperones in control and tumor tissues as well as cell lines was investigated and indeed, a significant upregulation of both the subunits of CAF1 - p60 and p150 and downregulation of HIRA (Fig 4A(i) and (ii)), was observed, very similar to the changes seen for their respective H3 variants. To further understand their correlation, the histone variants and their respective chaperones were profiled in various rat normal tissues. Indeed, as hypothesized in most of the tissues their expression pattern was correlated, wherever, elevated levels of H3.3 was observed, its chaperone HIRA was also found to be increased (Fig S5). This suggests that the simultaneous upregulation of the corresponding chaperone is important for the proper incorporation of the variant. An earlier report has shown that in the absence of its specific chaperone, the centromere-specific histone variant CENP-A is mis-incorporated into other genomic loci leading to chromosomal instability [27].

**Fig 4:**
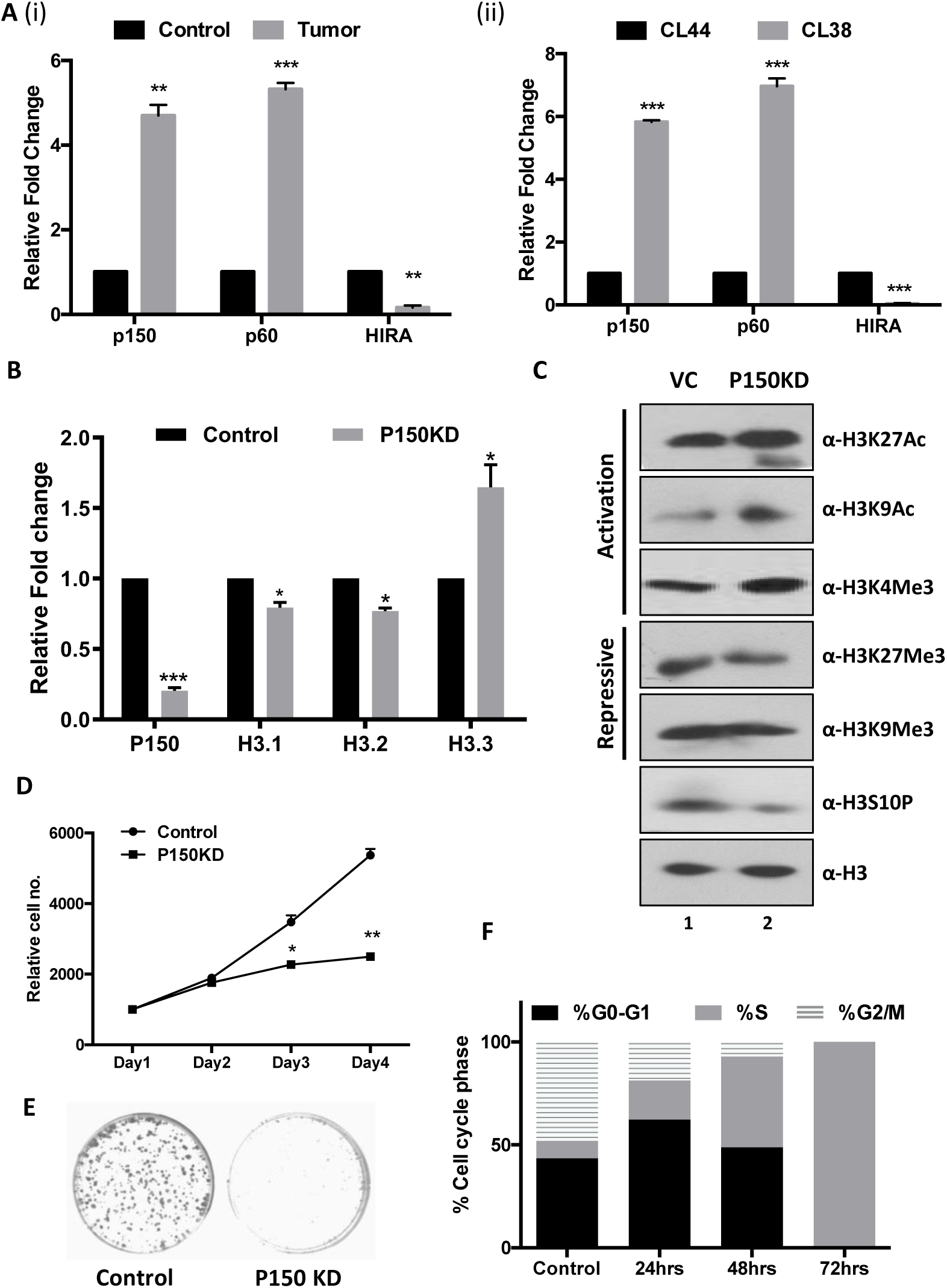
CAF1 suppression affects cell proliferation by cell cycle arrest in S-phase. (a) Quantitative real-time PCR data showing the relative expression levels of histone chaperones CAF-1 (P150 and P60) and HIRA in tissues (i) and cell lines (ii). (b) Relative transcript levels pf P150 and histone H3 variants – H3.1, H3.2 and H3.3 in knockdown and control cell lines. (c) Global histone PTM profiling for active and repressive marks by western blotting. Measurement of cell proliferation by (d) MTT assay and (e) Clonogenic assay. (f) Flow cytometry analysis at various time points of doxycycline induced P150 knockdown in comparison to control cells.

We next sought to understand the effect of the inducible knockdown of P150, a subunit of CAF-1, which is the histone chaperone of H3.2 and H3.1. The P150 levels were massively depleted at both the transcript and the protein levels post 48 hours of induction (Fig 4B and S6). The levels of histone transcripts post 48 hrs of doxycycline induction P150 knockdown was also measured (Fig 4B). Probing for various histone marks in control and P150 knockdown cell lines revealed an increase in the activation and a decrease in the repressive marks (Fig 4C). Further, MTT and clonogenic assay revealed a significant decrease in cell proliferation potential of the cells (Fig 4D and E), which also is reflected in the level of H3S10P, a mark correlated with the mitotic status of cells (Fig 4C). To understand the role of P150 depletion on cell proliferation, cell cycle status of the knockdown cells was assessed at various time points of knockdown. A gradual increase in the number of cells in the S-phase with the downregulation of p150, with 100% cells in the S-phase just after 72 hrs of doxycycline treatment (Fig 4F) was observed. This suggests that the changes observed upon P150 knockdown are probably due to the arrest of cells in S-phase and may not be attributed to the increase in the H3.3 levels.

### 3.5. H3.3 suppression recapitulates the tumor phenotype

To validate that the observed changes in cancer cells are indeed due to the loss of H3.3, an shRNA approach was employed for achieving H3.3 knockdown in the pre-neoplastic CL44 cells. The knockdown of H3.3 was validated at both the transcript (Fig 5A) and the protein level (Fig 5B). Interestingly, elevated expression of H3.2 and H3.1 was also seen upon the decrease in H3.3 expression. A significant increase in cell proliferation was observed post H3.3 loss as seen in MTT and clonogenic assays (Fig 5C, D). Western blotting with a panel of activation and repressive histone PTM marks revealed an increase in the repressive marks upon H3.3 loss (Fig 5E). H3.3 knockdown in hepatocytes led to an elevated expression of H3.2 and H3.1, probably for the maintenance of total H3 levels. The changes in the PTM marks could be attributed to both, H3.3 decrease and H3.2 increase. Intriguingly, an increased H3S10P and a drop in H4K16Ac-a hallmark of cancer accompany H3.3 knockdown, thus recapitulating the tumor specific changes. This allows us to conclude that the DNA methylation mediated loss of H3.3 leads to elevated cell proliferation and contributes to the carcinogenesis phenomenon. Of note, the observation in the drop of H4K16Ac upon knockdown of H3.3 incites that may be one of the mechanisms, the hallmark of cancer-loss of H4K16Ac is brought about might be via H3.3 loss along with hMOF dysregulation.

**Fig 5:**
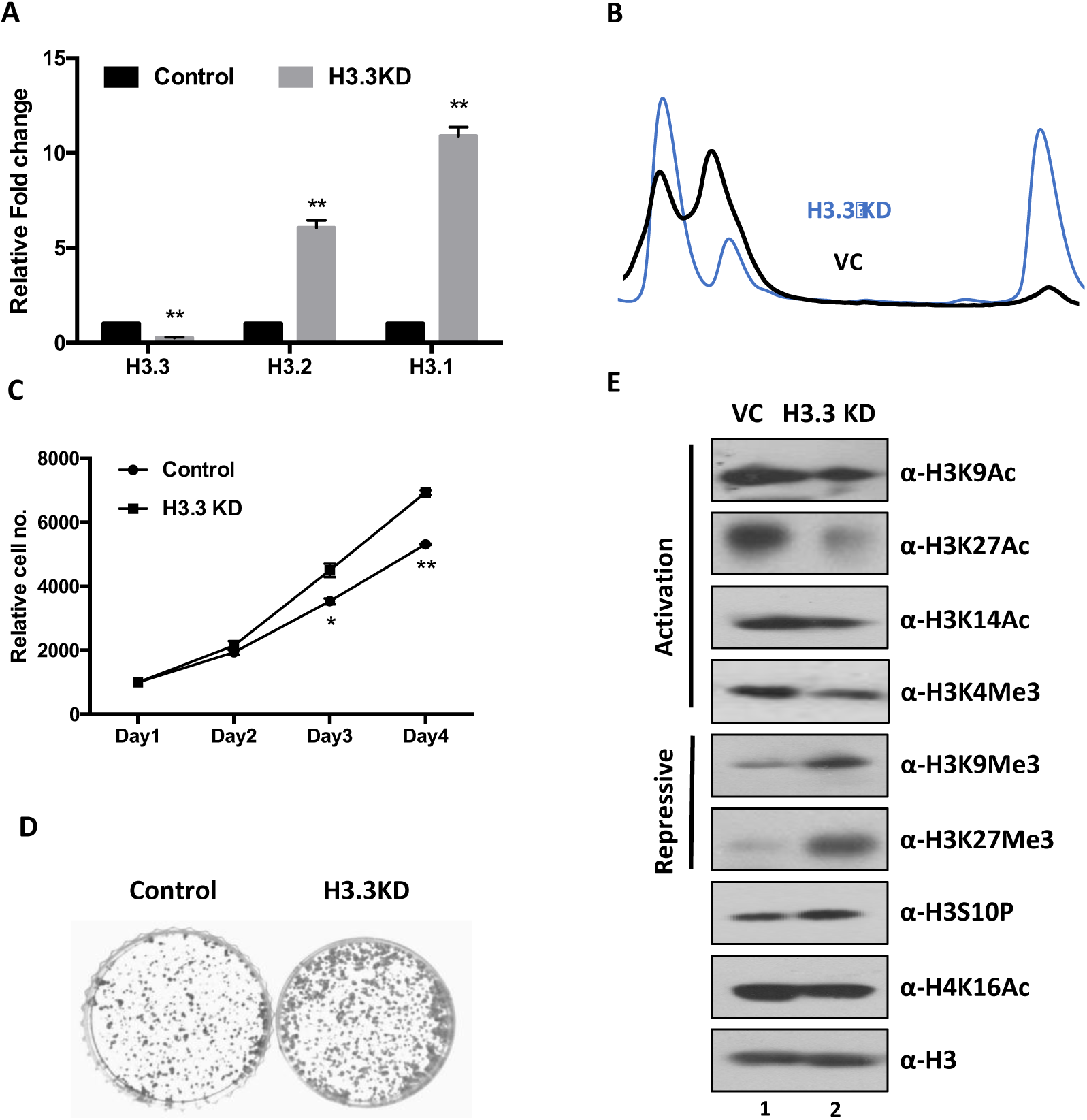
H3.3 Knockdown increases cell proliferation. (a) Quantitative real-time PCR data showing the relative expression levels of histone variants H3.3, H3.2 and H3.1 upon knockdown of H3.3 (b) RP-HPLC peak profile depicting the effect of H3.3 knock on H3 variant levels. Measurement of cell proliferation by (c) MTT assay and (d) Clonogenic assay. (e) Global histone PTM profiling for active and repressive marks by western blotting.

### 3.6. H3.3/ H3.2 levels regulate the expression of tumor suppressor genes

To understand the function of H3.3 in tumor cell proliferation, chromatin immunoprecipitation of MYC tagged H3.3 and H3.2 was performed in the CL44 cell line to check their relative enrichment on the various tumor suppressor genes. H3.3 seems to be associated with some tumor suppressors, unlike H3.2, suggesting that the deposition of H3.3 on these genes may govern their expression (Fig 6A). Further, the knockdown of H3.3 led to its loss on tumor suppressor genes and the simultaneous gain of H3.2 (Fig 6B).

**Fig 6:**
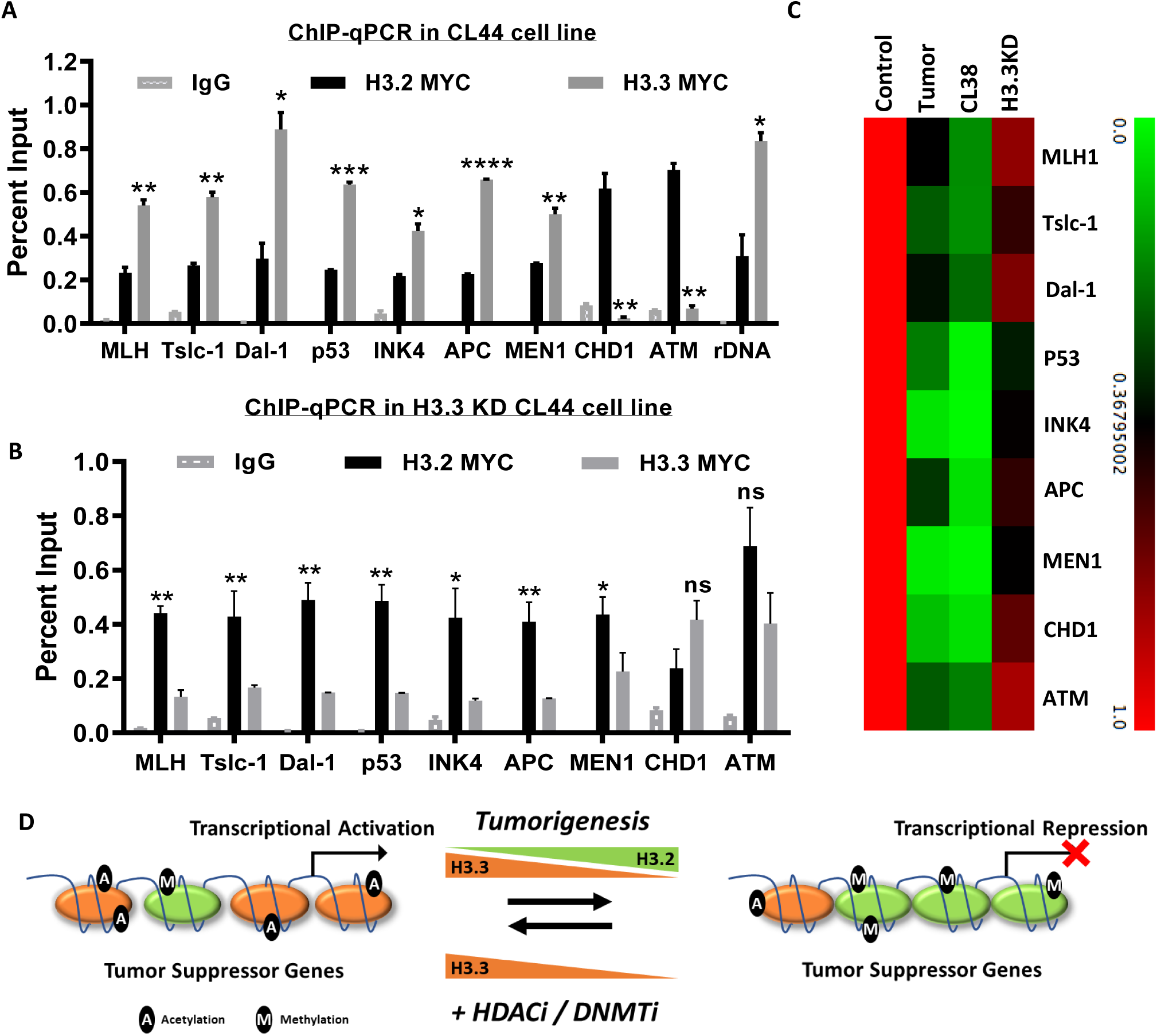
H3.3 governs tumor suppressor gene expression. (a) Measurement of relative enrichment of H3.2 and H3.3 on various tumor suppressor genes by ChIP-qPCR. rDNA locus was used as a control. IgG as an Isotype control. (b) ChIP-qPCR data showing relative levels of H3.3 and H3.2 on tumor suppressor genes upon H3.3 knockdown. rDNA locus was used as a positive control for the experiment, as it is mostly transcriptionally active (c) Relative transcript levels of tumor suppressors in tumor tissue, CL38 and H3.3 knockdown cell line normalized to GAPDH. (d) Dynamic changes in histone H3 variant profile along with their specific PTM’s influence tumor suppressor gene expression, which can be reversed by using DNMTi and HDACi.

To further validate that H3.3 levels indeed might be correlated to the expression status of these genes, their expression was monitored in tumor tissue, CL38 and H3.3 knockdown CL44 cells. In comparison to control, the tumor tissue and CL38 showed a tumor suppressor gene repression (Fig 6C). H3.3 knockdown led to a decrease in the gene expression of the monitored tumor suppressor genes, thus, providing a proof of principle that H3.3 loss aids in tumor cell phenotype acquirement. However, we cannot undermine the fact that an increase in H3.2 on these genes might also be responsible for their loss of expression. Interestingly, not all tumor suppressor genes are affected upon H3.3 knockdown, suggesting that probably there might be only a specific subset of genes that are under the control of H3.3 directly. Possibly, the elevated cell proliferation upon H3.3 loss may be due to loss of expression of APC, a negative regulator of the WNT signaling pathway.

## 4. Discussion

The epigenetic regulatory network has become a matter of intense investigation over the years from the perspective of disease pathology because unlike the genetic changes they are more amenable to reversal and are believed to be better targets for therapeutic intervention. In the present study, we have looked at the changes specifically in the histone H3 variants and their PTM profile in the HCC rat model system. We report a decreased expression of H3.3, along with an increase in H3.2 in HCC. In the past we had shown how DNA methylation is intimately linked to histone gene expression for H2A variants [28]. Here we demonstrate similar transcriptional regulation of H3 variants. Treatment with small molecule inhibitors, 5-Aza-C (DNMTi) and TSA (HDACi) led to the reversal of these changes (Fig 6D). Though previous studies have investigated independently the histone modification profile or the histone variant changes in cancer, however, to the best of our knowledge, our study for the first time shows that both the histone variant and the modification profile might be intimately linked. Association of H3.3 with transcriptional activation is a well-established fact, though, whether it is due to the PTMs it undergoes or due to its nature of forming inherently unstable nucleosome is not yet established.

Histone H3 mutations in cancer are well appreciated and know to effect global chromatin organization and thus gene expression. Not only the mutations but dysregulation of H3.3 expression in cancer has also been documented previously. H3.3 overexpression has been reported to promote lung cancer progression by regulating the expression of a key gene ‘GPR87’ involved in metastasis [11]. The MLL5 mediated loss of H3.3 in adult glioma has been attributed to be a trademark of cells undergoing differentiation [12]. Previous studies although have reported the alteration in histone variant profile, changes in their chaperones have not been investigated. Interestingly, the p150 subunit of the CAF1 chaperone is upregulated in many cancers and its elevated expression level was significantly correlated with poor clinicopathological features in patients with HCC and as an independent prognostic factor for predicting both the overall and disease-free 5-year survival [29]. p150 knockdown has been linked to a decrease in cell proliferation [30], and our results indicate that this can be due to the arrest of cells in S-phase probably owing to the decrease in the deposition of H3 variants needed for DNA compaction post their synthesis. The CAF-1 complex is also known to be important to safeguard somatic cell identity, as CAF-1 suppression led to a more accessible chromatin structure at enhancer elements early during reprogramming [31]. Though it is not yet understood if it is due to its capacity to deposit H3.2/H3.1 or not. Recently a study also highlighted the importance of the ERK signaling mediated, CAF1 suppression, and increased H3.3 deposition in governing the levels of pro aggressive markers during metastasis in breast cancer [31]. However, unlike this report, our study suggests that in our HCC model system there is a HIRA mediated decrease in H3.3, on tumor suppressor genes facilitates an increased cell proliferation and thus promoting cancer progression. This suggests a bimodal function of H3.3 in a context-dependent manner, probably during tumor initiation and progression as it regulates tumor suppressor gene expression and during metastasis, it governs expression of EMT markers. Such a kind of role has been already established for H3.3 during cell reprogramming. It plays an essential role in maintaining fibroblast identity during the initial phase of reprogramming, but its role is reversed at the later stages where it mediates the acquirement of pluripotency [31]. However, further detailed study during various stages of cancers is essential to establish such a context-dependent function of H3.3.

In summary, large scale chromatin structure changes occur as cells transition from normal to cancer state. These are brought about possibly by DNA methylation mediated downregulation of the histone variant H3.3 and upregulation of H3.2. These changes favor gene repression of various tumor suppressor genes, probably contributing to an elevated cell proliferation. Observations from our study prompt us to speculate that probably the reversal of such changes can be bought about using DNMT and HDAC inhibitors. Although there is much more to learn about how DNA methylation mediated changes in various histone variants and their modifications are initiated during carcinogenesis, these chromatin-mediated changes seen in our study provide clues to events initiating at the very earliest stages of neoplasia. Therefore, we conclude that the dysregulation of H3 variants acts as the central node of this reversible epigenetic process seen in cancer and therefore can have potential therapeutic implications.

## 5. Acknowledgments

Authors thank Dr. H. M. Rabes (University of Munich, Germany) for providing CL44 and CL38 cell lines. Also, the authors are grateful to all past and present members of Gupta Lab, ACTREC for valuable discussions. D.R. and S.B. were supported by CSIR fellowship. This work was supported in part by TMC-Intramural Research Grant and Department of Biotechnology, Government of India.

## Notes

The authors declare no potential conflicts of interest

### Competing Interest Statement

The authors have declared no competing interest.

## References

[1] M. V Brock, J.G. Herman, S.B. Baylin, Cancer as a manifestation of aberrant chromatin structure, Cancer J. 13 (2007).

[2] R.T. Kamakaka, S. Biggins, Histone variants: deviants?, Genes Dev. 19 (2005) 295–316. https://doi.org/10.1101/gad.1272805.

[3] A. Kapoor, M.S. Goldberg, L.K. Cumberland, K. Ratnakumar, M.F. Segura, P.O. Emanuel, S. Menendez, C. Vardabasso, G. Leroy, C.I. Vidal, D. Polsky, I. Osman, B.A. Garcia, E. Hernando, E. Bernstein, melanoma progression through regulation of CDK8, Nature. 468 (2010) 1105–1109. https://doi.org/10.1038/nature09590.

[4] A. Svotelis, N. Gévry, G. Grondin, L. Gaudreau, H2A.Z overexpression promotes cellular proliferation of breast cancer cells., Cell Cycle. 9 (2010) 364–70. https://doi.org/10.4161/cc.9.2.10465.

[5] D. Dryhurst, B. McMullen, L. Fazli, P.S. Rennie, J. Ausió, Histone H2A.Z prepares the prostate specific antigen (PSA) gene for androgen receptor-mediated transcription and is upregulated in a model of prostate cancer progression, Cancer Lett. 315 (2012) 38–47. https://doi.org/https://doi.org/10.1016/j.canlet.2011.10.003.

[6] D. Reddy, S. Gupta, Histone variant H3.3 and its future prospects in cancer clinic, J. Radiat. Cancer Res. 8 (2017) 77–81. https://doi.org/10.4103/jrcr.jrcr_4_17.

[7] C.-J. Lin, M. Conti, M. Ramalho-Santos, Histone variant H3.3 maintains a decondensed chromatin state essential for mouse preimplantation development, Development. 140 (2013) 3624–3634. https://doi.org/10.1242/dev.095513.

[8] B.T.K. Yuen, K.M. Bush, B.L. Barrilleaux, R. Cotterman, Histone H3. 3 regulates dynamic chromatin states during spermatogenesis, (2014) 3483–3494. https://doi.org/10.1242/dev.106450.

[9] H.T. Fang, C.A. El Farran, Q.R. Xing, L.F. Zhang, H. Li, B. Lim, Y.H. Loh, Global H3.3 dynamic deposition defines its bimodal role in cell fate transition, Nat. Commun. 9 (2018). https://doi.org/10.1038/s41467-018-03904-7.

[10] T. Meshi, K. Taoka, M. Iwabuchi, Regulation of histone gene expression during the cell cycle, Plant Mol. Biol. 43 (2000) 643–657. https://doi.org/10.1023/A:1006421821964.

[11] S. Park, E. Choi, M. Bae, S. Kim, J.B. Park, H. Yoo, J.K. Choi, Y. Kim, S. Lee, I. Kim, migration through intronic regulation, Nat. Commun. 7 (2016) 1–14. https://doi.org/10.1038/ncomms12914.

[12] R. Chromatin, M. Gallo, F.J. Coutinho, R.J. Vanner, D.P. Bazett-jones, M. Lupien, P.B. Dirks, M. Gallo, F.J. Coutinho, R.J. Vanner, T. Gayden, S.C. Mack, A. Murison, MLL5 Orchestrates a Cancer Self-Renewal State by Repressing the Histone Variant H3. 3 and Globally Article MLL5 Orchestrates a Cancer Self-Renewal State by Repressing the Histone Variant H3. 3 and Globally Reorganizing Chromatin, Cancer Cell. 28 (2015) 715–729. https://doi.org/10.1016/j.ccell.2015.10.005.

[13] S. Hua, C.B. Kallen, R. Dhar, M.T. Baquero, C.E. Mason, B.A. Russell, P.K. Shah, J. Liu, A. Khramtsov, M.S. Tretiakova, Genomic analysis of estrogen cascade reveals histone variant H2A. Z associated with breast cancer progression, Mol. Syst. Biol. 4 (2008) 188.

[14] J.C. Sporn, G. Kustatscher, T. Hothorn, M. Collado, M. Serrano, T. Muley, P. Schnabel, A.G. Ladurner, Histone macroH2A isoforms predict the risk of lung cancer recurrence, Oncogene. 28 (2009) 3423–3428.

[15] A. Ivashkevich, C.E. Redon, A.J. Nakamura, R.F. Martin, O.A. Martin, Use of the γ-H2AX assay to monitor DNA damage and repair in translational cancer research, Cancer Lett. 327 (2012) 123–133. https://doi.org/10.1016/j.canlet.2011.12.025.

[16] S.P. Khare, A. Sharma, K.K. Deodhar, S. Gupta, Brief Communication Overexpression of histone variant H2A. 1 and cellular transformation are related in N-nitrosodiethylamine-induced sequential hepatocarcinogenesis, (2011) 30–35.

[17] D. Shechter, H.L. Dormann, C.D. Allis, S.B. Hake, Extraction, purification and analysis of histones, Nat. Protoc. 2 (2007) 1445–1457. http://dx.doi.org/10.1038/nprot.2007.202.

[18] K.S. Pramod, K. Bharat, G. Sanjay, Mass spectrometry-compatible silver staining of histones resolved on acetic acid-urea-Triton PAGE, Proteomics. 9 (2009) 2589–2592.

[19] B.U. Holecek, R. Kerler, H.M. Rabes, Chromosomal Analysis of a Diethylnitrosamine-induced Tumorigenic and a Nontumorigenic Rat Liver Cell Line, Cancer Res. (1989) 3024–3028.

[20] N. Heintz, The regulation of histone gene expression during the cell cycle, 1088 (1991) 327–339.

[21] Y. Yamamoto, Y. Nishikawa, T. Tokairin, Y. Omori, K. Enomoto, Increased expression of H19 non-coding mRNA follows hepatocyte proliferation in the rat and mouse., J. Hepatol. 40 (2004) 808–14. https://doi.org/10.1016/j.jhep.2004.01.022.

[22] T. Tsujiuchi, E. Sugata, T. Masaoka, M. Onishi, H. Fujii, K. Shimizu, K. Honoki, Expression and DNA methylation patterns of Tslc1 and Dal-1 genes in hepatocellular carcinomas induced by N-nitrosodiethylamine in rats., Cancer Sci. 98 (2007) 943–8. https://doi.org/10.1111/j.1349-7006.2007.00480.x.

[23] H. Manoharan, K. Babcock, H.C. Pitot, Changes in the DNA methylation profile of the rat H19 gene upstream region during development and transgenic hepatocarcinogenesis and its role in the imprinted transcriptional regulation of the H19 gene, Mol. Carcinog. 41 (2004) 1–16. https://doi.org/10.1002/mc.20036.

[24] G.P. Pfeifer, Defining Driver DNA Methylation Changes in Human Cancer, Int. J. Mol. Sci. 19 (2018) 1166. https://doi.org/10.3390/ijms19041166.

[25] E. McKittrick, P.R. Gafken, K. Ahmad, S. Henikoff, Histone H3.3 is enriched in covalent modifications associated with active chromatin., Proc. Natl. Acad. Sci. (Of United States Am. 101 (2004) 1525–1530. https://doi.org/10.1073/pnas.0308092100.

[26] A. Hamiche, M. Shuaib, Chaperoning the histone H3 family, Biochim. Biophys. Acta - Gene Regul. Mech. 1819 (2012) 230–237. https://doi.org/10.1016/j.bbagrm.2011.08.009.

[27] T. Tomonaga, K. Matsushita, S. Yamaguchi, T. Oohashi, H. Shimada, T. Ochiai, K. Yoda, F. Nomura, Overexpression and mistargeting of centromere protein-A in human primary colorectal cancer., Cancer Res. 63 (2003) 3511–3516.

[28] M. Tyagi, D. Reddy, S. Gupta, Genomic characterization and dynamic methylation of promoter facilitates transcriptional regulation of H2A variants, H2A.1 and H2A.2 in various pathophysiological states of hepatocyte, Int. J. Biochem. Cell Biol. 85 (2017) 15–24. https://doi.org/10.1016/j.biocel.2017.01.019.

[29] Y. Wang, Chromatin assembly factor 1, subunit A (P150) facilitates cell proliferation in human hepatocellular carcinoma, (2016) 4023–4035.

[30] M. Hoek, B. Stillman, Chromatin assembly factor 1 is essential and couples chromatin assembly to DNA replication in vivo., Proc. Natl. Acad. Sci. U. S. A. 100 (2003) 12183–8. https://doi.org/10.1073/pnas.1635158100.

[31] S. Cheloufi, U. Elling, B. Hopfgartner, Y.L. Jung, J. Murn, M. Ninova, M. Hubmann, A.I. Badeaux, C. Euong Ang, D. Tenen, D.J. Wesche, N. Abazova, M. Hogue, N. Tasdemir, J. Brumbaugh, P. Rathert, J. Jude, F. Ferrari, A. Blanco, M. Fellner, D. Wenzel, M. Zinner, S.E. Vidal, O. Bell, M. Stadtfeld, H.Y. Chang, G. Almouzni, S.W. Lowe, J. Rinn, M. Wernig, A. Aravin, Y. Shi, P.J. Park, J.M. Penninger, J. Zuber, K. Hochedlinger, The histone chaperone CAF-1 safeguards somatic cell identity, Nature. 528 (2015) 218–224. https://doi.org/10.1038/nature15749.

