## Supplementary data for "DNA methylation mediated downregulation of histone H3 variant H3.3 affects cell proliferation contributing to the development of HCC"

### Supplementary 1

#### H3.2 Peptides

| Amino acid (Start-End) | Observed | Mol.Wt. (Expt) | Mol.Wt. (Calc) | ppm | Peptide |
|---|---|---|---|---|---|
| 28-41 | 1449.8177 | 1488.8105 | 1448.8212 | -7.20 | KSAPSTGGVKKPHR |
| 42-54 | 1600.9406 | 1599.9333 | 1599.9321 | -6.93 | YRPGTVALREIRR |
| 58-64 | 831.4884 | 830.4811 | 830.4861 | -6.05 | STELLIR |
| 65-73 | 1156.7266 | 1155.7193 | 1155.7240 | -4.11 | KLPFQRLVR |
| 71-84 | 1703.9254 | 1702.9181 | 1702.9366 | -10.8 | LVREIAQDFKTDLR |
| 85-117 | 3493.9232 | 3492.9154 | 3492.7984 | 32.7 | FQSAAIGALQEASEAYLVGLFEDT NLCAIHAKR |
| 118-130 | 1540.8909 | 1539.8836 | 1539.89.19 | -5.40 | VTIMPKDIQLARR |

#### H3.3 Peptides

| Amino acid (Start-End) | Observed | Mol.Wt. (Expt) | Mol.Wt. (Calc) | ppm | Peptide |
|---|---|---|---|---|---|
| 28-41 | 1433.8263 | 1432.8190 | 1432.8263 | -5.08 | KSAPATGGVKKPHR |
| 42-50 | 1032.5952 | 1031.5879 | 1031.5876 | -4.05 | YRPGTVALR |
| 58-64 | 831.4884 | 830.4811 | 830.4861 | -6.05 | STELLIR |
| 65-73 | 1156.7186 | 1155.7113 | 1155.7240 | -11 | KLPFQRLVR |
| 74-84 | 1335.6888 | 1334.6816 | 1334.6830 | -1.08 | EIAQDFKTDLR |
| 85-117 | 3513.7474 | 3512.7454 | 3512.7484 | 32.7 | FQSSAVMALQEASEAYLVGLFEDTNLCAI HAKR |
| 118-130 | 1540.8884 | 1539.8811 | 1539.8919 | -7.02 | VTIMPKDIQLARR |

Figure S1: Peptides identified for H3.2 and H3.3 by MS

### Supplementary 2

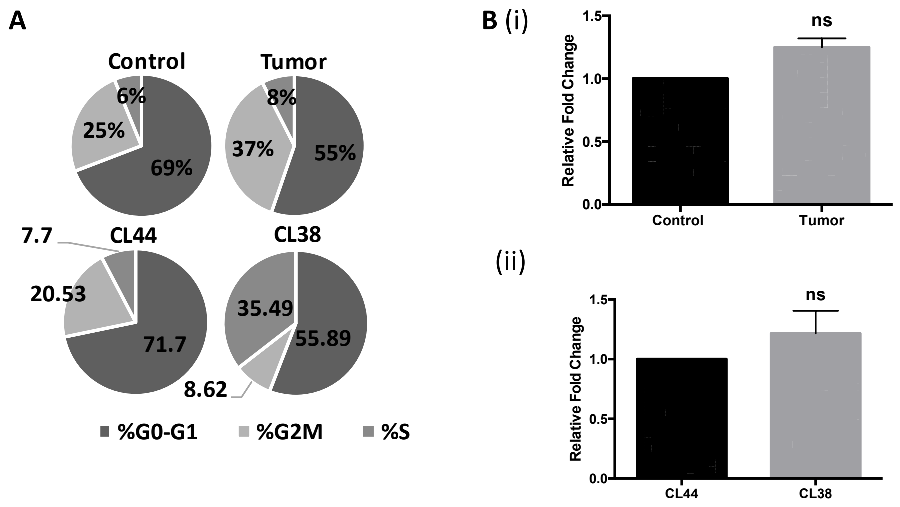


(A) Flow cytometry analysis depicting the percentage of cells in various phases of cell cycle in form of a pie diagram for both tissues and cell lines. (B) Monitoring the changes in H3.1 expression in (i) tissues and (ii) cell lines at transcript level.

### Supplementary 3

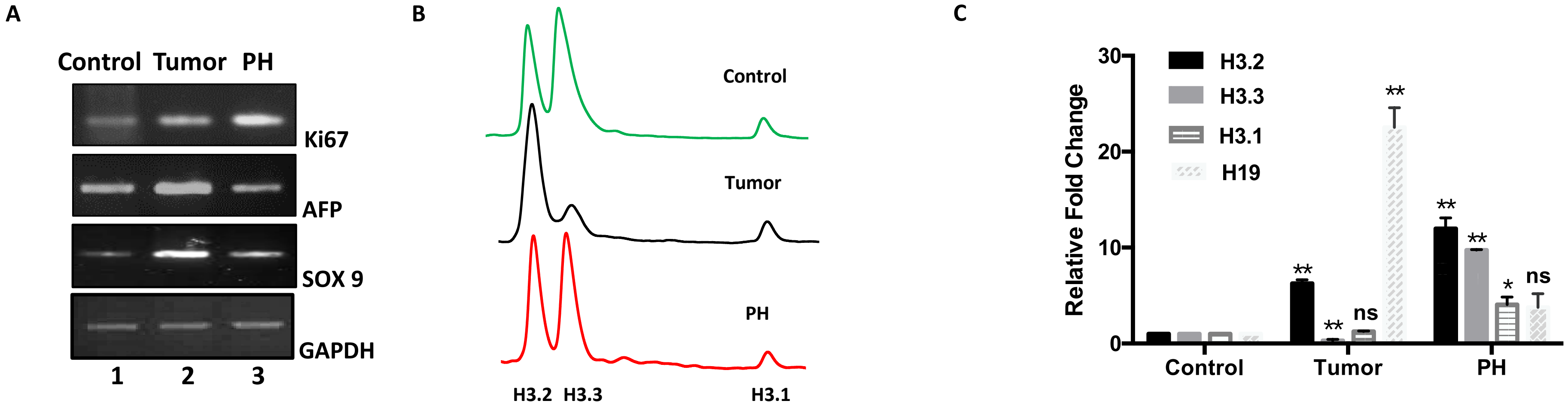


(A) Semi RT-PCR fortissues. GAPDH was used as a loading control. (B)RP-HPLC graph showing H3 variant peak profiles of control, tumor, partial hepatectomized (PH) liver tissues. (C) Quantitative PCR analysis of transcript levels of H3 variants in control, tumor and PH tissues. H19 was used as a control in the study, expression of which is known to change in the various pathophysiological states of liver.

### Supplementary 4

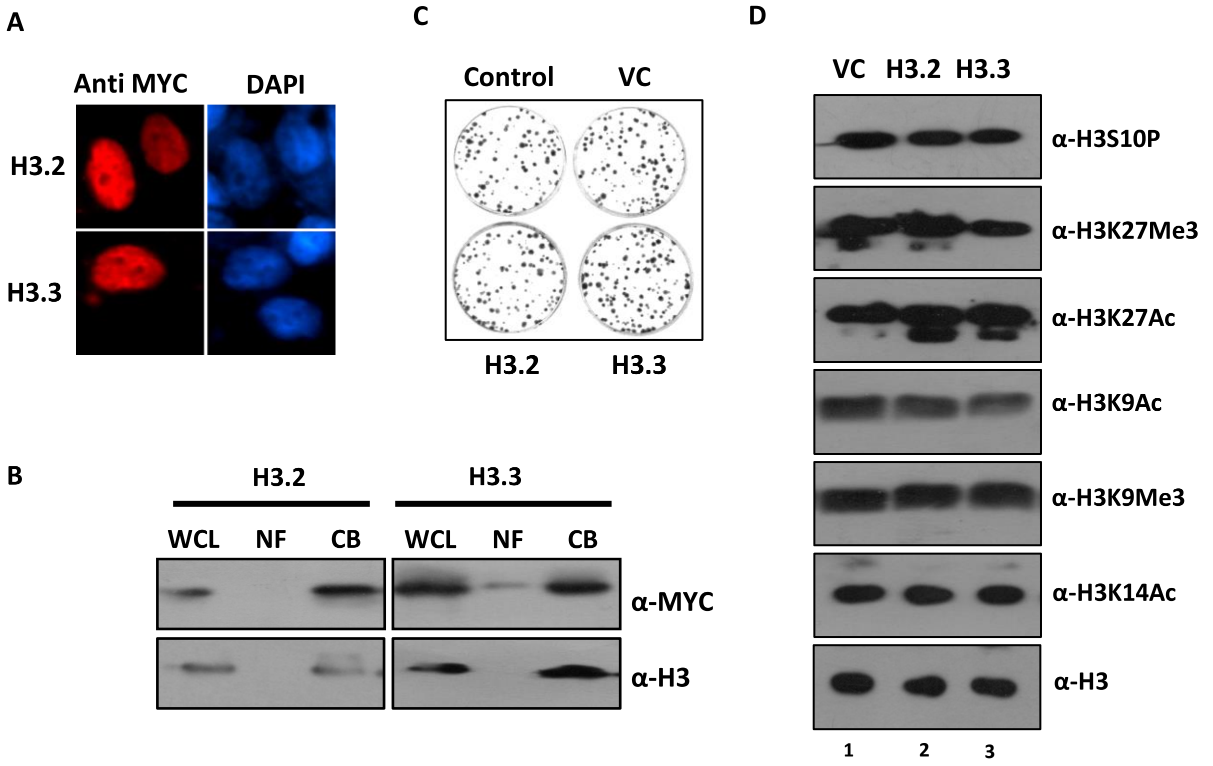


(A) Analysis of localization of MYC tagged H3.2 and H3.3 by Immunofluorescence microscopy. Red depicts the Alexa 568 staining. DAPI in blue counter stains the nuclei. (B) Cellular fractionation followed by immunoblotting with the marked antibodies to determine sub cellular distribution of histones. H3 western was used as the control. WCL- Whole Cell Lysate, NF- Nucleosolic Fraction and CF- Chromatin Fraction. (C) Colony formation assay obtained after plating 1000 cells and allowing growth for 14 days. The colonies were fixed and stained with crystal violet stain. (D) Western blotting of various histone PTM's.

Supplementary 5

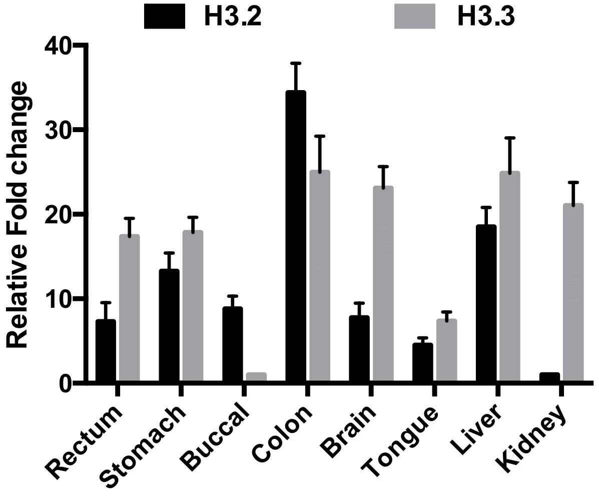


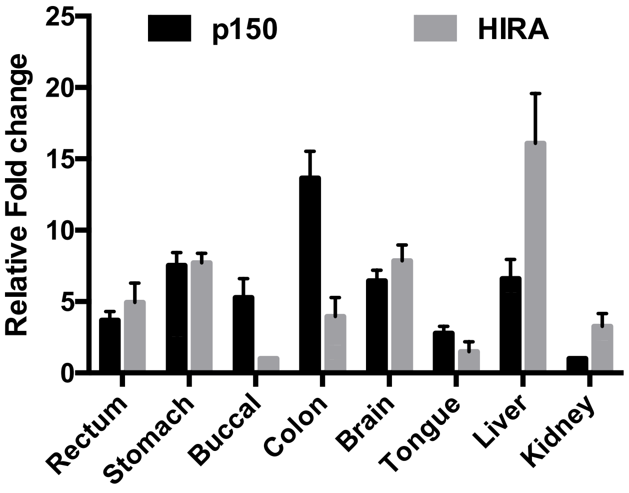


Graph depicting the relative levels of histone variants and histone chaperones transcripts across various rat normal tissues.

### Supplementary 6

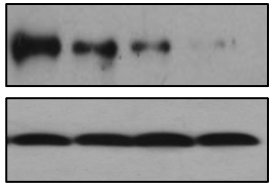


Doxycycline mediated depletion of P150 subunit of CAF-1 at various time points. Beta actin was used as a loading control. VC- Vehicle Control.
